# Diverse Lyme disease spirochete species evade restriction by human complement

**DOI:** 10.64898/2026.01.20.700503

**Authors:** Maryna Golovchenko, Lucie Krätzerová, Heather MacTavish, Vett Lloyd, Natalie Rudenko

**Affiliations:** Biology Centre CAS, Institute of Parasitology, 37005, Ceske Budejovice, Czech Republic; Military Health Institute, Biological Defence Department Techonin, 561 66, Techonin, CzechRepublic; Mt. Allison University, Sackville, New Brunswick, Canada E4L 1G7

**Keywords:** Borrelia, human complement, Lyme disease, spirochete mortality, infectivity, acute immune response, human biological sex

## Abstract

Disseminated human Lyme borreliosis (LB) is traditionally associated with invasive spirochete species from the *Borrelia burgdorferi* sensu lato (s.l.) complex. The *Borrelia burgdorferi* sensu lato complex consists of 23 spirochete species worldwide, with additional species being proposed. Interactions between the host immune system and *Borrelia* species have been studied for a diverse range of vertebrate reservoirs of LB spirochetes, including lizards and snakes, as well as the widely recognized rodent and bird reservoirs. Humans are the vertebrate species most susceptible to *B. burgdorferi* s.l. spirochetes, as demonstrated by the increasing number of diagnosed LB cases worldwide. However, few studies have evaluated differences in the borreliacidal action of human complement against specific *Borrelia* species. Using the serum sensitivity test, we analyzed whether complement-mediated killing of 10 spirochete species from the *B. burgdorferi* s.l. complex varies among healthy human individuals of different ages and sexes. Our results show that the 10 genospecies exhibit varying sensitivities to human complement and can be classified into three statistically distinct groups: high, medium, and low sensitivity. Complement sensitivity did not correlate with the known human health impact of these genospecies; similar resistance to the killing effects of human serum was found among *Borrelia* species that are major causes of LB worldwide and species with unclear pathogenicity to humans. Additionally, females showed reduced complement-mediated *Borrelia* killing compared to males for all *Borrelia* species in almost all age groups. Age and biological sex interacted for the *Borrelia* species most sensitive to human complement. Overall, the effectiveness of complement-mediated killing of the different *Borrelia* species tended to decrease with age, with more complex age-dependent changes observed for some *Borrelia* species.

## Introduction

The *Borrelia burgdorferi* sensu lato complex consists of 23 well-established and currently recognized spirochete species worldwide, along with two recently proposed South American species (1, 2). Five of these species, *B. afzelii*, *B. burgdorferi* sensu stricto (s.s.), *B. bavariensis, B. garinii*, and *B. mayonii*, are the major confirmed causes of Lyme borreliosis (LB), a multisystem disorder with a diverse spectrum of clinical manifestations. However, pathogenicity of other *Borrelia* species to humans has also been described (1–3). The *B. burgdorferi* s.l. complex has been divided into two groups: one includes *Borrelia* species clearly able to survive in humans after infection and that have been detected or isolated from humans, and so which are considered pathogenic. The other group includes those not yet detected in humans. However, the distinction between pathogenic and non-pathogenic *Borrelia* species may change as less well-studied species are further characterized. The pathogenic group includes 12 species: *B. afzelii, B. americana, B. andersonii, B. bavariensis, B. bissettii, B. burgdorferi* s.s.*, B. garinii, B. kurtenbachii, B. lusitaniae, B. mayonii, B. spielmanii* and *B. valaisiana* (1,3–5). The remaining 11 species have not yet been detected in humans and are therefore assumed to be non-pathogenic. These are *B. californiensis, B. carolinensis, B. chilensis, B. finlandensis, B. japonica, B. lanei, B. maritima, B. tanukii, B. turdi, B. sinica,* and *B. yangtzensis* (1,3–5). Among the species that have been detected in humans, the frequency of recovery varies greatly. While *B. afzelii, B. bavariensis, B. garinii, B. burgdorferi* s.s. or *B. mayonii* are commonly detected in diagnosed cases of LB in endemic regions of northern hemisphere (1), the number of patients in which *B. bissettii, B. kurtenbachii, B. spielmanii* or *B.valaisiana* have been detected is low, and detection of *B. americana, B. lusitaniae* or *B. andersonii* in humans is rare (2,6). What causes the unequal recovery of different *Borrelia* species from humans? Humans are accidental hosts for tick vectors, so exposure depends on a complex interplay of ecological and human behavioral factors. However, when a human is exposed to a specific spirochete species, what determines whether *Borrelia* can disseminate and cause disease in humans? Is the predominant detection of only a few *Borrelia* species due to biological constraints on the pathogenicity of each species, or is it the result of a historical research focus on *B. burgdorferi* s.s., *B. afzelii*, and *B. garinii* to the exclusion of other species?

Spirochetes first encounter the host immune system upon entering the skin, and subsequently in the interstitial fluid, bloodstream or lymphatic system as they travel to other tissues. When spirochetes contact complement in these tissues, the innate immune response is activated (8). Complement is an essential part of the innate immune system and consists of a series of proteins that, upon recognizing a pathogen, initiate a defensive cascade of interactions to lyse or engulf pathogens and to alert and enhance the adaptive immune response (7,8). At the beginning of infection, before pathogen-specific antibodies are generated, the invading spirochetes are attacked by the non-specific alternative complement pathway. *Borrelia* produces several Complement Regulator-Acquiring Surface Proteins, CRASPs, BbCRASP-1 to BbCRASP-5 that bind regulatory proteins of the alternative complement pathway, such as factor H and H-like protein 1 (FHL-1) (9–12). By binding these host regulatory proteins, spirochetes avoid recognition and eradication by the host complement system. It has been found that *Borrelia* host range is determined in both wild and laboratory animals, in part, by its sensitivity to complement of a particular vertebrate host species (13–21) as the ability to bind factor H and FHL-1 depends on the *Borrelia* genotype (11,15). As a result, this divides spirochete species into serum-sensitive and serum-resistant strains. Here, we determine whether this serum sensitivity underlies the pathogenicity of different *Borrelia* species for a specific host species, such as humans.

## Materials and Methods

### Cultivation of Borrelia species

Ten *Borrelia* species, *B. burgdorferi* s.s. B31, *B. afzelii* CB43, *B. garinii* PBi, *B. bissettiae* DN127, *B. kurtenbachii* 25015, *B. carolinensis* SCGT18, *B. americana* SCW30F, *B. bavariensis* 20047, *B. valaisiana* VS116 and *B. andersonii* 21038, were selected for serum sensitivity analysis. All spirochetes were kept as a frozen stocks at −80°C. Freshly prepared BSK-H medium (22,23) was inoculated with frozen low passage spirochete culture under sterile conditions. No antibiotics were used in spirochetes re-cultivation. Newly seeded cultures were incubated at +34°C with regular monitoring of the growth by dark-field microscopy as described earlier (24). The presence of a single *Borrelia* species in the culture was confirmed by PCR on genomic DNA isolated from a culture aliquot using the DNeasy Blood and Tissue kit (Qiagen, Germany) as described earlier with the following sequence of amplified fragments for re-identification of spirochete species (25). Cultures, prepared in this way were grown until density reached 10^7^ cells/ml and were then used in the serum sensitivity test.

### Human serum samples

To obtain the human samples, the office of a general practitioner physician, where patients’ blood was collected on a daily basis, was contacted. The blood was collected by medical professionals in the course of regular patient healthcare unrelated to this study. During the blood collection all patients were informed about the opportunity to provide residual blood samples (2 ml) for research and verbal consent was obtained; 148 blood samples were collected in this way; two donors declined participation. The blood samples provided to the research team were anonymous, with only age and biological sex provided for each sample, and no record connecting blood samples to patient identity was made. Ethical consent to conduct this study was provided by the Ethics Committee of the Biological Center in České Budějovice (5/2024) and the Mount Allison University Research Ethics Board (104186). Samples were divided into 6 age categories of donors: category 1 (21-30 years), 2 (31-40 years), 3 (41-50 years), 4 (51-60 years), 5 (61-70 years) and 6 (over 71 years). Each category was further divided into samples from i) biological females and ii) biological males. Following clinical use of the blood, the residual sample was brought to the lab. The blood samples were centrifuged at 2,000 rpm at room temperature for 15 minutes to separate blood cells and serum and then filtered through a 0.22 μm sterile PES syringe filter (Fisher Scientific, USA). The presence of anti-*Borrelia* antibodies was screened by the rapid immunochromatographic test (Lymetest (Lyme IgG/IgM rapid test for whole blood/serum/plasma), Biorepair, Germany). Ten positive samples (8 male and 2 female), all from people of age groups 3 and 4 (41-60 years), were excluded from the study as seropositive.

Complement sensitivity testing was conducted according to the protocol established by Kurtenbach and colleagues (14). Fifty microliters of *Borrelia* culture, for each species, cell density 10^7^ cells/ml, was added in ratio 1:1 to individual serum sample and co-incubated at 34°C for 24 hours in microtiter plates. Samples were then supplemented with 100µl of dilution buffer (2% BSA, 5.4mM glucose in sterile PBS) and transferred to 5 ml BD Falcon tubes. One µl of propidium-iodide (PI) was then added and reactions were incubated in the dark for 15 minutes after the addition of PI. The sensitivity of different *Borrelia* species to each group of human sera was analyzed by flow cytometry using measurements of 30,000 events, with clusters of cells digitally removed. The intensity of propidium iodide fluorescence was measured in the PE-Texas Red-A 616/23 channel. The resulting values measured using a flow cytometer indicate the percentage of dead *Borrelia* in the individual samples. The average percentage of dead *Borrelia* in each group was calculated from 10 measurements. Each time the assay was run, a live spirochete control (the same spirochete species in BSK-H medium) and a dead spirochete control (spirochete culture heated at 56°C for 30 minutes) were used. In total, 1,200 complement sensitivity tests were performed and all samples and controls were analyzed in the same way. For all analyzed species, the average mortality rate for all *Borrelia* species under these experimental conditions was below 10%. Supplemental Table 1 provides the raw data.

### Statistical analysis

The survival of *Borrelia* species (Supplemental Table 1) were first compared using a Kruskal-Wallis test, followed by Dunn’s post hoc pairwise comparisons (Supplemental Table 2), to categorize them into low, medium, and high sensitivity groups based on significantly different levels of complement-mediated killing. Once divided into sensitivity groups, the effects of age, biological sex, and their interaction were assessed using an Aligned Rank Transform (ART) ANOVA (Supplemental Table 3). Where significant interactions were detected, post hoc Wilcoxon rank sum tests were conducted within each age category to compare males and females separately, allowing assessment of sex-specific effects within each age category. All post hoc comparisons were corrected for multiple testing using the Holm-Bonferroni method. Patterns across age and sex combinations were visualized using Principal Coordinate Analysis (PCoA) based on Bray-Curtis distances.

## Results

### Human host complement-mediated Borrelia mortality is dependent on Borrelia species

The human complement system is an essential part of innate immunity and has antibacterial properties. Examining *Borrelia* cell viability after exposing different spirochete species to human complement indicates whether the complement is protective against that *Borrelia* species. Evaluation of the results of host complement killing of the different *Borrelia* species was assessed using the Kruskal-Wallis H test, using the combined age and gender data for the complement-mediated *Borrelia* mortality for each *Borrelia* species (Figure 1, Supplemental Table 1). A Kruskal-Wallis rank-sum test showed a significant difference in complement-mediated killing among the Borrelia species (χ²(9) = 914.12, p < 2.2 × 10⁻16), indicating a highly statistically significant difference in complement sensitivity between the species. The Dunn test was used to do post hoc pairwise comparisons. Of the 45 comparisons, only 9 were not significant (Bafz – Bam, Bafz – Bbav, Band – Bbiss, Bam – Bbss, Band – Bcar, Bbiss – Bcar, Bafz – Bkurt, Bcar – Bgar, Bbav – Bkurt; Figure 1).

**Figure 1.**
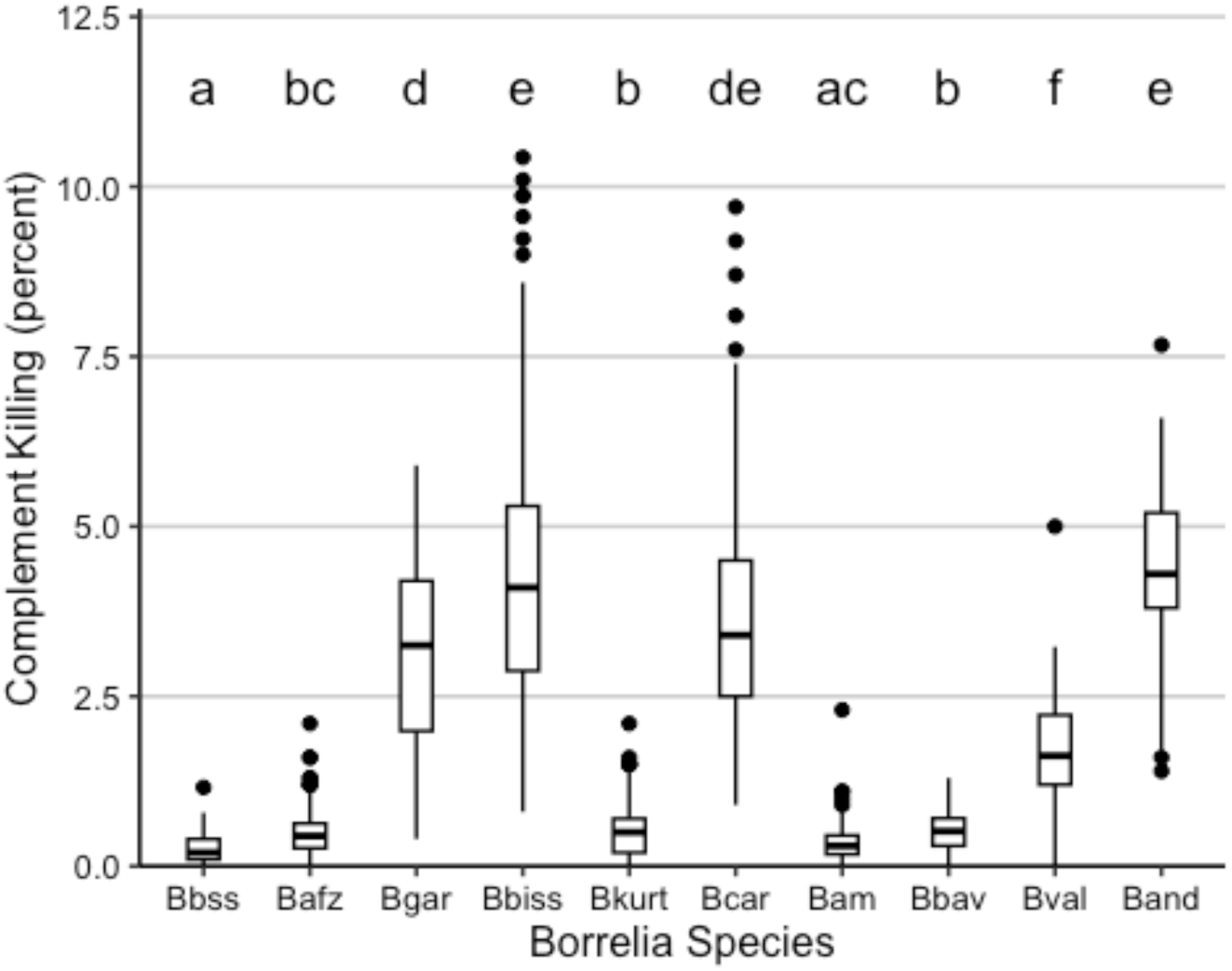
Percent complement-mediated killing of 10 different *Borrelia* species. Box plots sharing letters are not significantly different from in pairwise comparisons. For example, Bbss (a) is significantly different to Bafz (bc) but is not significantly different to Bam (ac). Bval (f) is significantly different to all of the other species. Bbss = B. burgdorferi sensu stricto, Bafz = B. afzelii, Bgar = B. garinii, Bbis = *B. bissettiae*, Bkurt = *B. kurtenbachii*, Bcar = *B. carolinensis*, Bam = *B. americana*, Bbav = *B. bavariensis*, Bval = *B. valaisiana*, Band = *B. andersonii*.

These findings delineated three groups with statistically significant differences in their complement sensitivity (Figure 1). The group with the lowest mortality when challenged by host complement was *B. burgdorferi* sensu stricto (Bbss, a), *B. afzelii* (Bafz, bc), *B. kurtenbachii* (Bkurt, b), *B. americana* (Bam, ac), and *B. bavariensis* (Bbav, b); *B. burgdorferi* s.s. and *B. americana* had the lowest mortality in this group. *B. afzelii, B. bavariensis* and *B. kurtenbachii* similarly shared low sensitivity suggesting that the infectious potential of *B. kurtenbachii* is comparable to that of the widely distributed and pathogenic *B. afzelii* and *B. bavariensis.* This may indicate that increased human infections with *B. kurtenbachii* can be expected in its distribution area. The sensitivity of *B. valaisiana* (f) to human serum was intermediate. The group with the greatest sensitivity to human complement included *B. garinii*, *B. bissettiae, B. carolinensis* and *B. andersonii,* which was the species most sensitive to human serum (Figure 1). The complement sensitivity for *B. bissettiae* and *B. carolinensis* were comparable to that of *B. garinii,* predicting a similar ability of these species to survive in humans after infection.

### Effect of age and biological sex on complement-mediated Borrelia mortality

To test the effect of age and sex on complement-mediated Borrelia killing, it was first necessary to assess whether there was an interaction between age and sex to determine if the results could be readily interpreted. If there was no interaction, the main effects could be interpreted directly; however, if an interaction was present, the results could not be interpreted without further analysis and consideration of limitations. Aligned Rank Transform ANOVA tests (Supplemental Table 3) demonstrated that the low (Bbss (a), Bafz (bc), Bkurt (b), Bam (ac), Bbav (b)) and medium sensitivity (Bval) groups showed no significant interaction between age and sex of the host (p = 0.63347 for the low sensitivity group and p = 0.1346905 for the medium sensitivity group). However, the high sensitivity group (Bgar (d), Bbiss (e), Bcar (de), Band) showed a significant interaction (p = 0.00080765), so the main effects of age or sex cannot be interpreted independently.

For the low and medium sensitivity groups there is a significant effect of age on complement-mediated *Borrelia* mortality (p = 1.02 × 10⁻^5^ and p = 0.0018468, respectively; Figure 2A and 2B). For *B. burgdorferi* s.s*, B. afzelii* and *B. americana*, the effect appears fairly linear and sensitivity declines with age. For *B. kurtenbachii*, the effect is opposite, with a seeming increase in complement-mediated mortality as age increases. *Borrelia bavariensis* and *B. valaisiana* show a pattern in which the highest complement-mediated *Borrelia* mortality occurs in the 51-60 year-old age group, with reduced *Borrelia* killing in both younger and older individuals. This pattern may also apply to most high sensitivity *Borrelia* species (Figure 2C), although the age and sex interaction makes the results harder to interpret.

**Figure 2.**
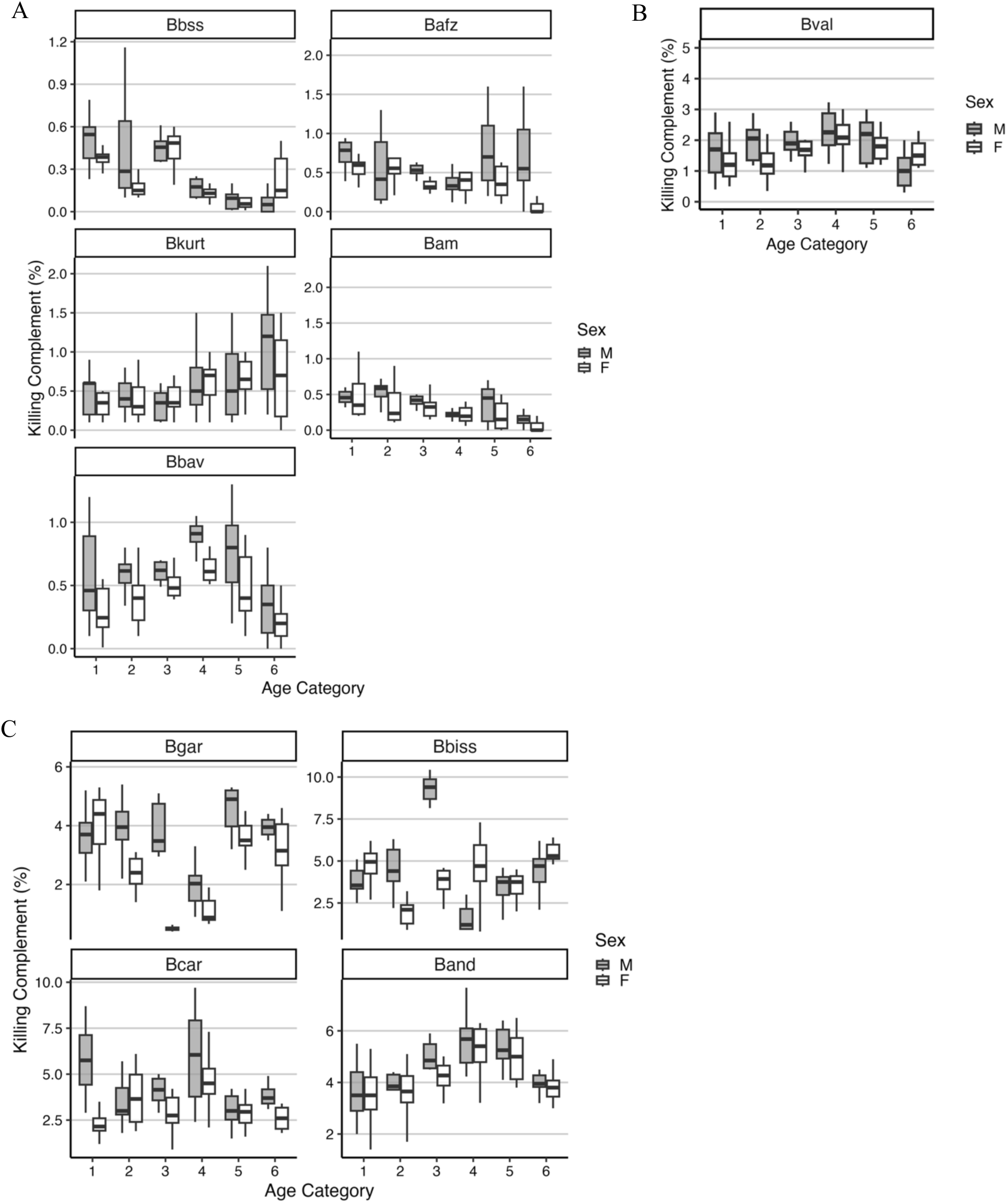
The effect of age and host biological sex on complement-mediated *Borrelia* lethality. The charts show the percent complement-mediated *Borrelia* lethality for age categories 1-6 where age category 1 = 21-30 years, 2 = 31-40 years, 3 = 41-50 years, 4 = 51-60 years, 5 = 61-70 years, 6 = 70+ years, with host biological sex indicated by box shading (dark = male, unfilled = femails). Bbss = B.burgdorferi sensu stricto, Bafz = B.afzelii, Bgar = B.garinii, Bbis = B. bissettiae, Bkurt = B.kurtenbachii, Bcar = B.carolinensis, Bam = B.americana, Bbav = B.bavariensis, Bval = B.valaisiana, Band = B.andersonii. A. Results for the low sensitivity *Borrelia* species. B. Results for the medium sensitivity *Borrelia* species. C. Results for the high sensitivity *Borrelia* species.

Biological sex consistently affected complement-mediated *Borrelia* killing, with females showing reduced killing compared to males across nearly all age groups and *Borrelia* species (Figure 2A, B, C). For the low and high sensitivity groups, the decreased *Borrelia* mortality in females was statistically significant (p = 2.62 × 10⁻^6^ for the low sensitivity group). For the single medium sensitivity species, the effect was not statistically significant (p = 0.1158315), although visual examination of Figure 2B shows decreased complement-mediated *Borrelia* killing in females in several age groups. For the high sensitivity group, the interaction between age and sex complicates interpretation of the effect of biological sex on complement-mediated *Borrelia* killing; however, this effect is again visually apparent. Pairwise post hoc comparisons, with age and sex combined into groups, using the Wilcoxon rank sum test with continuity correction (Supplemental Table 4), showed p values below 0.05 for males aged 41-50 years (group 3M). This is illustrated with a Principal Coordinate Analysis (Figure 3) showing the relationships among different age groups across all *Borrelia* species based on Bray-Curtis distances. The group of 41-to 50-year-old males (3M) appears as the most dissimilar, highlighting a possible synergistic effect of age and gender on complement-mediated *Borrelia* killing in this group. These findings collectively suggest that females may be more vulnerable to infection by *Borrelia* species.

**Figure 3.**
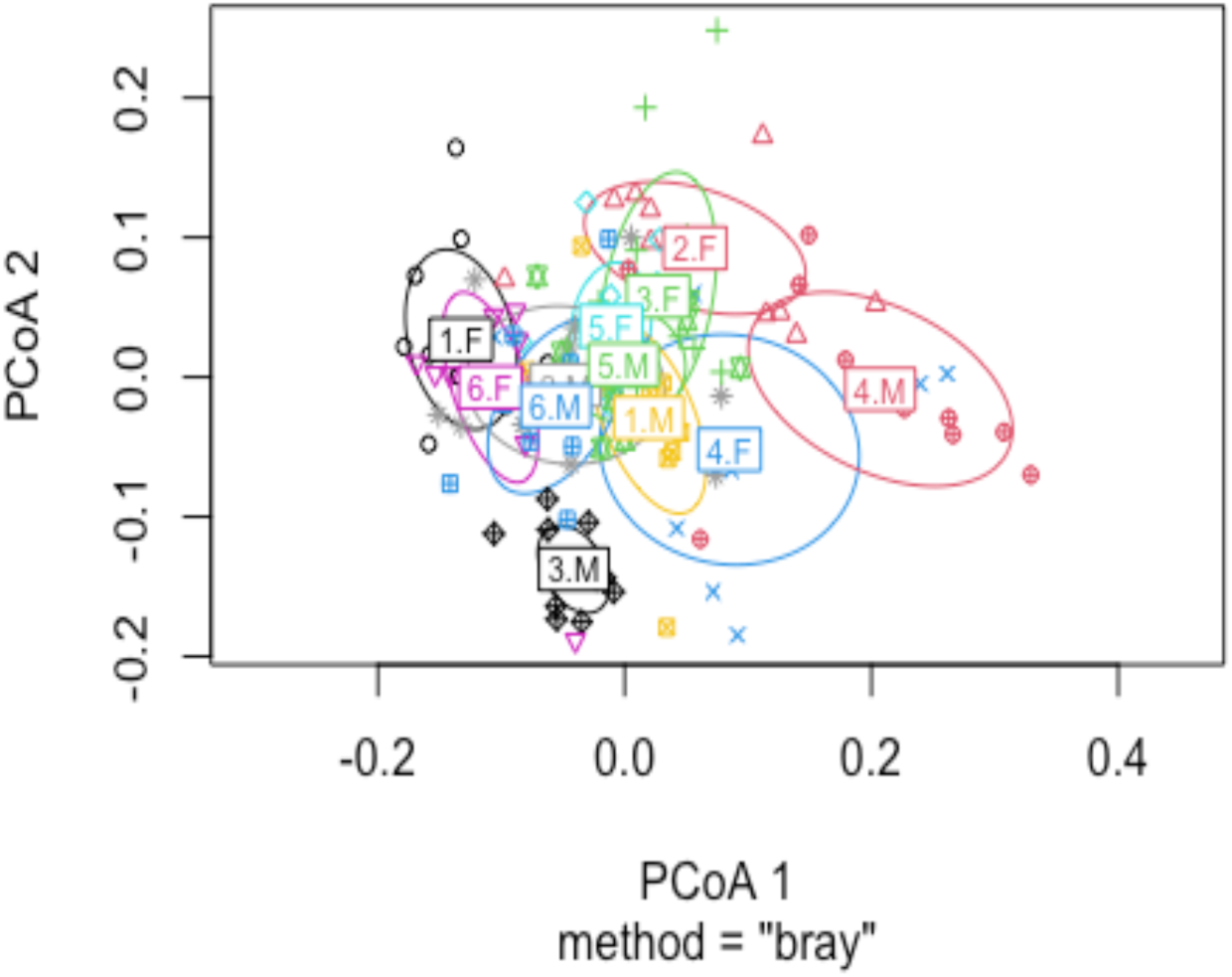
Principal Coordinate Analysis (PCoA) of complement-mediated *Borrelia* killing across all *Borrelia* species, based on Bray-Curtis distances. The plot illustrates relative dissimilarities among all age-sex combinations, with the 3M group appearing as the most distinct.

## Discussion

The findings of this study have several important implications. We found that resistance to human serum was similar between some well-known pathogenic species recognized as the major etiological agents of Lyme borreliosis worldwide and those less frequently reported in humans or with unclear pathogenicity toward humans. This suggests that the human complement would not serve as a barrier to infection by these other species should humans be exposed to them. Whether other factors protect against infection by these species, or whether infections by *Borrelia* species considered non-pathogenic or uncharacterized have occurred but been overlooked, is an important question that requires attention as climate change and other factors alter the distribution of ticks, their wild reservoir hosts, and *Borrelia* spirochetes.

For example, the resistance of *B. americana* to human serum was the same as that of *B. burgdorferi*, the major cause of Lyme disease in North America. This suggests that human infection could be expected in areas where *B. americana* is present in reservoir species and tick vectors, as documented in southeastern and western United States and western Canada (26–28). Similarly, complement-mediated mortality was not statistically different between *B. kurtenbachii* and the highly infective and pathogenic Eurasian species *B. afzelii* and *B. bavariensis*. Strains closely related to *B. kurtenbachii* have been detected in ticks and humans in Europe (30–32), the United States (33–35), and Canada (36). These data predict a risk of human infections in both Europe and North America. Likewise, *B. bissettiae* and *B. carolinensis* have a sensitivity to complement-mediated killing similar to that of *B. garinii*, the second most important cause of human Lyme borreliosis in Europe (37–39). Although *B. bissettiae* and *B. carolinensis* have previously been considered exclusively North American species, detection and isolation of *B. bissettiae* from humans have already been reported in both Europe and North America (37–39).

The results of this study also support the previous separation of *B. bavariensis* and *B. kurtenbachii* from their parental species. *B. bavariensis* was separated from *B. garinii* based on MLST and MSLA analyses (40). Similarly, *B. kurtenbachii* was separated from the *B. bissettiae* group, with which it had previously been grouped (41). The sensitivity of *B. garinii* and *B. bavariensis* to human serum complement differs, as is also the case for *B. bissettiae* and *B. kurtenbachii*, reinforcing the biological distinction between these species.

Collectively, the results presented here highlight the need for greater clinical awareness of infections caused by *Borrelia* species other than the commonly reported *B. burgdorferi* s.s., *B. garinii*, and *B. afzelii*. Our findings also confirm an urgent need for expanded wildlife and vector surveillance of diverse *Borrelia* species. The overall risk to human health depends on both exposure to diverse *Borrelia* species, which is determined by their presence in animal reservoir hosts and tick vectors, and the pathogenicity of the *Borrelia* species. In areas where these spirochetes are present in wildlife and ticks, our results indicate that the human complement system does not prevent human infection. Our finding of a largely uniform reduction in complement protection in females across all age groups against the tested *Borrelia* species has important public health implications. Interesting, with the caveat of a very small sample size reduced complement-induced spirochete mortality against different *ospC* variants of *B. burgdorferi* s.s. was also found by Pearson and colleagues (29). A final implication of these findings is that labeling a spirochete species as “non-infectious” to humans is provisional. Such a definition should include a deeper analysis of the specific complement receptors of each *Borrelia* species, rather than relying on the number of cases in which a particular *Borrelia* species has been identified in humans. Detection of *Borrelia* by standard two-tiered clinical serological testing is based on a limited number of antigens from a limited number of *Borrelia* species and would be expected to fail, and indeed has failed, to detect infections by unexpected species (30). Identification of complement receptors on spirochetes, which they use as a protective mechanism against host complement, represents one of the fascinating and critically important areas that need to be explored for a better understanding of the interactions between spirochetes and hosts.

## Acknowledgments

This research was partially funded by grant 25-17301S from Czech Science Foundation (CSF), grant LUC23151 INTER-COST from Ministry of Education, Youths and Sport of the CR (NR and MG) and the Natural Sciences and Engineering (NSERC) grant to VKL.

## Declaration of Interest Statement

No potential conflict of interest was reported by the authors.

**Maryna Golovchenko:** Conceptualization; Investigation; Methodology; Supervision; Validation: Writing-Original draft preparation.

**Lucie Ticha:** Formal analysis; Investigation; Methodology; Validation; Visualization.

**Heather MacTavish:** Methodology; Validation; Visualization; Statistics.

**Vett Lloyd:** Methodology; Supervision; Validation: Visualization; Writing-original draft preparation; Writing- Review and Editing.

**Natalie Rudenko:** Conceptualization; Funding Acquisition; Investigation; Methodology; Project administration; Resources; Supervision; Validation: Visualization; Writing-original draft preparation; Writing- Review and Editing

---

Supplementary Table 1. Raw complement-mediated killing (%) of 10 *Borrelia* species by human sera, separated by age category and biological sex.

Note: Bbss = B. burgdorferi sensu stricto; Bafz = B. afzelii; Bgar = B. garinii; Bbis = B. bissettiae; Bkurt = B. kurtenbachii; Bcar = B. carolinensis; Bam = B. americana; Bbav = B. bavvariensis; Bval = B. valaisiana; Band = B. andersonii. Values represent percentage mortality. Data are stratified by age category (1 (21–30), 2 (31–40), 3 (41–50), 4 (51–60), 5 (61–70), 6 (70+) and sex (F/M).

[Separate file]

**Supplementary Table 2.** Kruskal-Wallis rank sum test and post-hoc Dunn’s test results comparing complement-mediated killing across 10 *Borrelia* species. P-values adjusted using the Holm-Bonferroni method. Significance levels: * p ≤ 0.05, ** p ≤ 0.01, *** p ≤ 0.001. Kruskal-Wallis rank sum test: χ² = 914.12, df = 9, p < 2.2 × 10⁻^16^

| Comparison | Z score | Raw p-value | Adjusted p-value |
| --- | --- | --- | --- |
| Bafz - Bam | 2.27 | 0.0115 | 0.104 |
| Bafz - Band | -14.72 | $2.35 \times 10^{-49}$ | $9.16 \times 10^{-48}$ *** |
| Bam - Band | -16.99 | $4.55 \times 10^{-65}$ | $2 \times 10^{-63}$ *** |
| Bafz - Bbav | -0.64 | 0.26 | 0.78 |
| Bam - Bbav | -2.92 | 0.00177 | 0.0213* |
| Band - Bbav | 14.08 | $2.59 \times 10^{-45}$ | $9.32 \times 10^{-44}$ *** |
| Bafz - Bbis | -13.68 | $6.45 \times 10^{-43}$ | $2.26 \times 10^{-41}$ *** |
| Bam - Bbis | -15.96 | $1.31 \times 10^{-57}$ | $5.37 \times 10^{-56}$ *** |
| Band - Bbis | 1.04 | 0.149 | 0.897 |
| Bbav - Bbis | -13.04 | $3.65 \times 10^{-39}$ | $1.17 \times 10^{-37}$ *** |
| Bafz - Bbss | 3.28 | 0.000511 | 0.00664 ** |
| Bam - Bbss | 1.01 | 0.156 | 0.779 |
| Band - Bbss | 18.01 | $8.76 \times 10^{-73}$ | $3.94 \times 10^{-71}$ *** |
| Bbav - Bbss | 3.93 | $4.29 \times 10^{-5}$ | 0.000729 *** |
| Bbis - Bbss | 16.97 | $7.2 \times 10^{-65}$ | $3.09 \times 10^{-63}$ *** |
| Bafz - Bcar | -12.78 | $1 \times 10^{-37}$ | $3.11 \times 10^{-36}$ *** |
| Bam - Bcar | -15.06 | $1.56 \times 10^{-51}$ | $6.22 \times 10^{-50}$ *** |
| Band - Bcar | 1.94 | 0.0264 | 0.211 |
| Bbav - Bcar | -12.14 | $3.2 \times 10^{-34}$ | $9.28 \times 10^{-33}$ *** |
| Bbis - Bcar | 0.9 | 0.185 | 0.738 |
| Bbss - Bcar | -16.07 | $2.11 \times 10^{-58}$ | $8.87 \times 10^{-57}$ *** |
| Bafz - Bgar | -11.03 | $1.43 \times 10^{-28}$ | $4 \times 10^{-27}$ |
| Bam - Bgar | -13.3 | $1.18 \times 10^{-40}$ | $3.88 \times 10^{-39}$ *** |
| Band - Bgar | 3.7 | 0.00011 | 0.00176 ** |
| Bbav - Bgar | -10.38 | $1.48 \times 10^{-25}$ | $3.71 \times 10^{-24}$ *** |
| Bbis - Bgar | 2.66 | 0.00395 | 0.0434 * |
| Bbss - Bgar | -14.31 | $9.39 \times 10^{-47}$ | $3.48 \times 10^{-45}$ *** |
| Bcar - Bgar | 1.76 | 0.0394 | 0.276 |
| Bafz - Bkurt | -0.3 | 0.38 | 0.38 |
| Bam - Bkurt | -2.58 | 0.00498 | 0.0498 * |
| Band - Bkurt | 14.42 | $2.04 \times 10^{-47}$ | $7.74 \times 10^{-46}$ *** |
| Bbav - Bkurt | 0.34 | 0.368 | 0.735 |
| Bbis - Bkurt | 13.38 | $4.08 \times 10^{-41}$ | $1.39 \times 10^{-39}$ *** |
| Bbss - Bkurt | -3.59 | 0.000166 | 0.00232** |
| Bcar - Bkurt | 12.48 | $4.83 \times 10^{-36}$ | $1.45 \times 10^{-34}$ *** |
| Bgar - Bkurt | 10.72 | $4.04 \times 10^{-27}$ | $1.09 \times 10^{-25}$ *** |
| Bafz - Bval | -7.36 | $9.08 \times 10^{-14}$ | $2.09 \times 10^{-12}$ *** |
| Bam - Bval | -9.63 | $2.86 \times 10^{-22}$ | $6.87 \times 10^{-21}$ *** |
| Band - Bval | 7.36 | $9.22 \times 10^{-14}$ | $2.03 \times 10^{-12}$ *** |
| Bbav - Bval | -6.72 | $9.18 \times 10^{-12}$ | $1.84 \times 10^{-10}$ *** |
| Bbis - Bval | 6.32 | $1.3 \times 10^{-10}$ | $2.47 \times 10^{-9}$ *** |
| Bbss - Bval | -10.65 | $9.08 \times 10^{-27}$ | $2.36 \times 10^{-25}$ *** |
| Bcar - Bval | 5.42 | $2.94 \times 10^{-8}$ | $5.29 \times 10^{-7}$ *** |
| Bgar - Bval | 3.66 | 0.000124 | 0.00186 ** |
| Bkurt - Bval | -7.06 | $8.52 \times 10^{-13}$ | $1.79 \times 10^{-11}$ *** |

**Supplementary Table 3.** Results of the ART ANOVAs, testing the effect of age category and sex on complement-mediated killing. Significance levels: * p ≤ 0.05, ** p ≤ 0.01, *** p ≤ 0.001.

| Effect | df | F-value | p-value |
| --- | --- | --- | --- |
| Age Category | 5 | 1.4656 | 0.19963670 |
| Sex | 1 | 19.1766 | $1.4715 \times 10^{-5}$ *** |
| Age Category × Sex | 5 | 4.2831 | 0.00080765 *** |

(3B) The medium-sensitivity *Borrelia* group.
| Effect | df | F-value | p-value |
| --- | --- | --- | --- |
| Age Category | 5 | 4.1158 | 0.0018468 ** |
| Sex | 1 | 2.5130 | 0.1158315 |
| Age Category × Sex | 5 | 1.7262 | 0.1346905 |

(3C) The low-sensitivity *Borrelia* group.
| Effect | df | F-value | p-value |
| --- | --- | --- | --- |
| Age Category | 5 | 6.31031 | $1.0202 \times 10^{-5}$ *** |
| Sex | 1 | 22.51654 | $2.6188 \times 10^{-6}$ *** |
| Age Category × Sex | 5 | 0.68695 | 0.63347 |

**Supplementary Table 4.** Wilcoxon rank sum test results comparing complement-mediated killing between sexes within each age category for high-, medium-, and low-sensitivity *Borrelia* groups. P-values adjusted using the Holm-Bonferroni method. Significance levels: * p ≤ 0.05, ** p ≤ 0.01, *** p ≤ 0.001.

| Sensitivity | Age Category | Raw p-value | Adjusted p-value | Higher Group |
| --- | --- | --- | --- | --- |
| High | 1 | 0.214 | 0.643 | M |
| High | 2 | 0.00254 | 0.0127 * | M |
| High | 3 | $1.57 \times 10^{-6}$ | $9.43 \times 10^{-6}$ *** | M |
| High | 4 | 0.962 | 0.962 | F |
| High | 5 | 0.143 | 0.573 | M |
| High | 6 | 0.376 | 0.751 | M |
| Medium | 1 | 0.364 | 1 | M |
| Medium | 2 | 0.0233 | 0.14 | M |
| Medium | 3 | 0.307 | 1 | M |
| Medium | 4 | 0.52 | 1 | M |
| Medium | 5 | 0.595 | 1 | M |
| Medium | 6 | 0.111 | 0.555 | F |
| Low | 1 | 0.00831 | 0.0499 * | M |
| Low | 2 | 0.0344 | 0.103 | M |
| Low | 3 | 0.0143 | 0.0716 | M |
| Low | 4 | 0.493 | 0.493 | F |
| Low | 5 | 0.075 | 0.15 | M |
| Low | 6 | 0.0255 | 0.102 | M |

